# ATGL-dependent white adipose tissue lipolysis controls hepatocyte PPARα activity

**DOI:** 10.1101/2021.01.28.428684

**Authors:** Anne Fougerat, Gabriele Schoiswohl, Arnaud Polizzi, Marion Régnier, Carina Wagner, Sarra Smati, Tiffany Fougeray, Yannick Lippi, Frederic Lasserre, Valentine Melin, Blandine Tramunt, Chantal Alkhoury, Anthony Emile, Michael Schupp, Pierre Gourdy, Patricia Dubot, Thierry Levade, Sandrine Ellero-Simatos, Laurence Gamet-Payrastre, Ganna Panasyuk, Ez-Zoubir Amri, Catherine Postic, Walter Wahli, Nicolas Loiseau, Alexandra Montagner, Dominique Langin, Achim Lass, Hervé Guillou

**Author notes:** **Corresponding authors**: Achim Lass –, Institute of Molecular Biosciences, NAWI Graz, University of Graz, Heinrichstraße 31/II, A-8010 Graz, Austria; BioTechMed-Graz, Austria. Hervé Guillou -, Toxalim (Research Center in Food Toxicology), INRAE, ENVT, INP- PURPAN, UMR 1331, UPS, Université de Toulouse, Toulouse, France. Equal contributions (Achim Lass, Hervé Guillou).

## Abstract

**Objective:** In hepatocytes, peroxisome proliferator-activated receptor α (PPARα) acts as a lipid sensor that regulates hepatic lipid catabolism during fasting and orchestrates a genomic response required for whole-body homeostasis. This includes the biosynthesis of ketone bodies and the secretion of the starvation hormone fibroblast growth factor 21 (FGF21). Several lines of evidence suggest that adipose tissue lipolysis contributes to this specific process. However, whether adipose tissue lipolysis is a dominant signal for the extensive remodeling of liver gene expression dependent on PPARα has not been investigated.

**Methods:** First, using mice lacking adipose tissue lipolysis through adipocyte-specific deletion of adipose triglyceride lipase (ATGL), we characterized the responses dependent on adipocyte ATGL during fasting. Next, we performed liver whole genome expression analysis in fasted mice upon deletion of adipocyte ATGL or hepatocyte PPARα. Finally, we tested the consequences of hepatocyte-specific PPARα deficiency during pharmacological induction of adipocyte lipolysis with a β_3_-adrenergic receptor agonist.

**Results:** In the absence of ATGL in adipocytes, ketone body and FGF21 productions were impaired in response to starvation. Liver transcriptome analysis revealed that adipocyte ATGL is critical for regulation of hepatic gene expression during fasting and highlighted a strong enrichment in PPARα target genes in this condition. Genome expression analysis confirmed that a large set of fasting-induced genes are sensitive to both ATGL and PPARα. Adipose tissue lipolysis induced by acute activation of the β_3_-adrenergic receptor also triggered PPARα-dependent responses in the liver, supporting a role for adipocyte-derived fatty acids as dominant signals for hepatocyte PPARα activity. In addition, the absence of hepatocyte PPARα altered brown adipose tissue (BAT) morphology and reduced UCP1 expression upon stimulation of the β_3_-adrenergic receptor. In agreement with this finding, mice lacking hepatocyte PPARα showed decreased tolerance to acute cold exposure.

**Conclusions:** These results underscore the central role of hepatocyte PPARα in the sensing of adipocyte-derived fatty acids and reveal that its activity is essential for full activation of BAT. Intact PPARα activity in hepatocytes is required for cross-talk between adipose tissues and the liver during fat mobilization during fasting and cold exposure.

## 1. Introduction

The liver is one of the key organs involved in the control of energy homeostasis in response to nutrient availability. It can rapidly switch from glucose utilization and lipogenesis in the fed state to glucose production, fatty acid oxidation and ketogenesis during starvation. These processes strongly rely on transcriptional regulation. In response to nutrients and hormones, several specific transcription factors are activated and regulate genes involved in metabolic pathways in order to maintain energy homeostasis [1]. Among the transcription factors highly active in the liver, peroxisome proliferator-activated receptor α (PPARα) is critical for the adaptive response to fasting [2–5]. PPARα is a ligand-activated transcription factor that belongs to the nuclear receptor superfamily and is a member of the peroxisome proliferator-activated receptor (PPAR) family, which also includes PPARβ/δ and PPARγ. Endogenous ligands of PPARα include fatty acids and various fatty acid-derived compounds [6]. In hepatocytes, PPARα controls the expression of several genes involved in whole-body fatty acid homeostasis, including fatty acid degradation and ketogenesis, allowing the liver to catabolize fatty acids for its own utilization and to provide energy substrates to other organs during food deprivation [2–4]. Consequently, mice with hepatocyte-specific deletion of *Pparα* have impaired ketone body production in response to fasting, and develop spontaneous hepatic steatosis during aging [2]. In addition, PPARα is required for the expression of fibroblast growth factor 21 (FGF21) [7–9], a hepatokine with systemic metabolic effects and hepatoprotective properties [10,11].

During fasting, adipocyte lipolysis in white adipose tissue (WAT) releases a large amount of fatty acids that are delivered to the liver. Fatty acids derived from adipose tissue are essential for fasting-induced expression of hepatic FGF21 [12]. In addition to its role in the mobilization of stored triglycerides, recent studies reported that lipolysis in WAT, but not in BAT, is required to fuel thermogenesis during fasting [13,14] and in response to cold exposure [15,16]. Fatty acids released from WAT during lipolysis stimulate the activation of the transcription factor HNF4α in the liver and serve as substrates for the hepatic production of acylcarnitines to fuel BAT thermogenesis during cold exposure [17].

Since PPARα acts as a lipid sensor in the liver and circulating free fatty acids increase during fasting, it has been suggested that fatty acids released from white adipocytes during fasting act as PPARα ligands to increase its activity in the liver. However, the contribution of adipose tissue lipolysis to hepatocyte PPARα-dependent gene regulation has not been studied. Here, we used mice models of defective adipocyte lipolysis (*Atgl*^adipo−/−^) and hepatocyte PPARα (*Pparα*^hep−/−^) to provide evidence that adipose-derived fatty acids represent a prominent signal for PPARα activity in the liver. In addition, we report that hepatocyte PPARα is required for BAT thermogenesis. Overall, these results identify white adipose ATGL-dependent lipolysis as a key regulator of hepatic function through the control of gene expression and hepatocyte PPARα as a major player of the cross-talk between adipose tissues and the liver during ATGL-dependent lipolysis.

## 2. Material and methods

### 2.1. Mouse models

*Pparα* hepatocyte-specific knockout (*Pparα*^hep−/−^) mice were generated at INRAE’s rodent facility (Toulouse, France) by mating the floxed-*Pparα* mouse strain with C57BL/6J albumin-Cre transgenic mice, as described previously [2], to obtain albumin-Cre^+/−^*Pparα*^flox/flox^ mice. Albumin-Cre^−/−^*Pparα*^flox/flox^ *(Pparα*^hep+/+^) littermates were used as controls. Genotyping was performed using an established protocol [2].

Adipocyte-specific *Atgl* knockout (*Atgl*^adipo−/−^) mice were bred at the animal facility of the University of Graz. They were originally generated at the animal facility of the University of Pittsburgh by mating the floxed-*Atgl* mouse strain with adiponectin-Cre mice, as described previously [18], to obtain adiponectin-Cre^+/−^*Atgl*^flox/flox^ mice. Adiponectin-Cre^−/−^*Atgl*^flox/flox^ mice *(Atgl*^adipo+/+^) were used as controls.

### 2.2. *In vivo* experiments

*In vivo* studies were performed in compliance with the European guidelines for the use and care of laboratory animals, and approved by an independent ethics committee under the authorization numbers 14005-2018030917086471 and 17430-2018110611093660, and by the Austrian Federal Ministry for Science, Research, and Economy (protocol number GZ: 39/9-4/75 ex 2017/18). All mice were housed at 21– 23 °C on a 12-hour light (ZT0–ZT12) / 12-hour dark (ZT12–ZT24) cycle and had free access to the standard rodent diet (Safe 04 U8220G10R) and tap water. ZT stands for Zeitgeber time; ZT0 is defined as the time when the lights are turned on. All mice used in this study were males and were sacrificed at ZT16.

#### 2.2.1. Fasting experiment and β_3_-adrenergic receptor activation in the adipocyte *Atgl*-deficient mouse model

Six 12-week-old mice were fed *ad libitum* or fasted for 24 h (starting at ZT16). For β_3_-adrenergic receptor activation, 12 week-old mice were fed *ad libitum* and given CL316243 (3 mg/kg body weight; Sigma Aldrich) or vehicle (0.5% carboxymethylcellulose in sterilized water) by gavage at ZT10 and sacrificed at ZT16 (n = 6/genotype/experimental condition).

#### 2.2.2. β_3_-adrenergic receptor activation and cold exposure in the hepatocyte *Pparα*-deficient mouse model

Twelve week-old mice were transferred in a ventilated cabinet at the specific temperature of 30 °C (thermoneutrality) 2 weeks before experiments. Mice were fed *ad libitum* or fasted at ZT0 and given CL316243 (3 mg/kg body weight; Sigma Aldrich) or vehicle (0.5% carboxymethylcellulose in sterilized water) by gavage at ZT10 and sacrificed at ZT16 (n = 6–10/genotype/experimental condition). For cold exposure, mice were single-housed without nesting material and had free access to food and water. Mice were transferred to 4 °C for 5 hours. Body temperature was taken every 15 minutes during the first hour, every 30 minutes during the second hour, then every hour with a RET-3 rectal probe (Physitemp) using a digital thermometer (Bioseb) (n = 8/genotype).

### 2.3. Blood and tissue sampling

Prior to sacrifice, blood was collected into EDTA-coated tubes (Sarstedt, K3E tubes) from the submandibular vein. Plasma was prepared by centrifugation (1500 g, 15 min, 4 °C) and stored at −80 °C. Following sacrifice by cervical dissociation, the liver and the BAT were removed, weighed, dissected, and prepared for histology, or snap frozen in liquid nitrogen and stored at −80 °C until use.

### 2.4. Blood glucose and plasma analysis

Free fatty acids (FFAs) were determined from plasma samples using a COBASMIRA+ biochemical analyzer (Anexplo facility, Toulouse, France). Plasma insulin concentration was measured using the Insulin Mouse Serum Assay HTRF Kit (Cisbio) (WE-MET in Toulouse). Blood glucose was measured with an Accu-Chek Guide glucometer (Roche Diagnostics). β-Hydroxybutyrate was measured with Optium β-ketone test strips that carried Optium Xceed sensors (Abbott Diabetes Care). Plasma FGF21 was assayed using the rat/mouse FGF21 ELISA kit (Sigma) according to the manufacturer’s instructions.

Free carnitine and acylcarnitines were measured from plasma (10 μl) spotted on filter membranes (Protein Saver 903 cards; Whatman), dried, and then treated as reported [19]. Briefly, acylcarnitines were derivatized to their butyl esters and treated with the reagents of the NeoGram MSMS-AAAC kit (PerkinElmer). Their analysis was carried out on a Waters 2795/Quattro Micro AP liquid chromatography–tandem mass spectrometer (Waters, Milford, MA).

### 2.5. Gene expression

Total cellular RNA was extracted from liver and BAT samples using TRIzol reagent (Invitrogen). RNA was quantified using a NanoDrop (Nanophotometer N60, Implen). Two micrograms of total RNA were reverse transcribed using the High-Capacity cDNA Reverse Transcription Kit (Applied Biosystems) for real-time quantitative polymerase chain reaction (qPCR) analyses. Primers for Sybr Green assays are presented in **Supplementary Table S1**. Amplifications were performed on an ABI Prism 7300 Real-Time PCR System (Applied Biosystems). qPCR data were normalized to the level of TATA-box binding protein (TBP) messenger RNA (mRNA) and analyzed with LinRegPCR (v2017.1) to derive mean efficiency (N0) [20,21].

Transcriptome profiles were obtained for 6 liver samples per group at the GeT-TRiX facility (GénoToul, Génopole Toulouse Midi-Pyrénées) using Sureprint G3 Mouse GE v2 microarrays (8×60K, design 074809, Agilent Technologies), according to the manufacturer's instructions. For each sample, Cyanine-3 (Cy3) labeled cRNA was prepared from 200 ng of total RNA using the One-Color Quick Amp Labeling kit (Agilent Technologies), according to the manufacturer's instructions, followed by Agencourt RNAClean XP (Agencourt Bioscience Corporation, Beverly, Massachusetts). Dye incorporation and cRNA yield were determined using a Dropsense 96 UV/VIS droplet reader (Trinean, Belgium). Next, 600 ng of Cy3-labeled cRNA were hybridized on the microarray slides, following the manufacturer’s instructions. Immediately after washing, slides were scanned on an Agilent G2505C Microarray Scanner using Agilent Scan Control A.8.5.1 software and the fluorescence signal extracted using Agilent Feature Extraction software v10.10.1.1 with default parameters. Microarray data and experimental details are available in NCBI's Gene Expression Omnibus (GEO) database (accession numbers GSE165699 and GSE165558).

### 2.6. Histology

Paraformaldehyde-fixed, paraffin-embedded BAT was sliced into 3-μm sections and stained with hematoxylin and eosin (H&E).

### 2.7. UCP1 protein expression

Ten milligrams of BAT was homogenized in RIPA buffer (50 mM Tris-HCl, pH 7.4, 150 mM NaCl, 2 mM EDTA, 0.1% SDS, 1 mM PMSF, 1% NP40, 0.25% sodium deoxycholate, proteinase and phosphatase inhibitors) and centrifuged 30 min at 13,000 g. The protein concentration in supernatants was measured using a Pierce BCA Protein Assay Kit (Thermo Scientific). Proteins were denatured in Laemmli buffer (62.5 mM Tris HCl, pH 6.8, 2% SDS, 10% glycerol, 5% β-mercaptoethanol, 0.02% bromophenol blue), separated by SDS-polyacrylamide gel electrophoresis (12%), and transferred onto nitrocellulose membrane. The membrane was blocked with 5% BSA in TBST (10 mM Tris-HCl, 150 mM NaCl, 0.05% Tween-20). Immunodetection was performed using anti-UCP1 (1:1,000, Abcam, ab10983) and anti-Hsp90 (1:1,000, Cell Signaling, #4877) overnight at 4 °C, followed by horseradish peroxidase–conjugated secondary antibody (anti-rabbit 1:5,000) for 1 h at room temperature. Signals were acquired using enhanced chemiluminescence using Clarity Western ECL Substrate (Bio-Rad) and a ChemiDoc Touch Imaging System (Bio-Rad).

### 2.8. Statistical analyses

Statistical analyses on biochemical and qPCR data were performed using GraphPad Prism for Windows (version 7.00; GraphPad Software). Two-way ANOVA was performed, followed by appropriate post-hoc tests (Sidak’s multiple comparisons test) when differences were found to be significant (*p* < 0.05). When only 2 groups were compared, the Student *t*-test was used; *p* < 0.05 was considered significant.

Microarray data were analyzed using R [22] and Bioconductor packages [23] as described in GEO accession numbers GSE165699 and GSE165558. Raw data (median signal intensity) were filtered, log2 transformed, corrected for batch effects (microarray washing bath and labeling serials), and normalized using the qsmooth method [24]. A model was fitted using the limma lmFit function [25]. Pair-wise comparisons between biological conditions were applied using specific contrasts. A correction for multiple testing was applied using the Benjamini-Hochberg procedure [26] to control the false discovery rate (FDR). Probes with an FDR ≤0.05 were considered to be differentially expressed between conditions.

Gene ontology and transcription factor enrichment analysis were performed using Metascape [27]. Gene network analysis was performed using String (version 11.0) [28].

## 3. Results

### 3.1. Adipose ATGL-dependent lipolysis is required for fasting-induced transcriptional responses in the liver

In order to investigate the specific contribution of WAT lipolysis to hepatic lipid homeostasis, we used mice with defective lipolysis due to adipocyte-specific deletion of *Atgl* through the use of the adiponectin promoter for Cre-mediated *Atgl* gene deletion [18]. To investigate the liver response to fasting, wild-type (*Atgl*^adipo+/+^) and adipocyte-specific *Atgl* knockout mice (*Atgl*^adipo−/−^) were either fed *ad libitum* or fasted for 24 h. As expected, adipocyte *Atgl* deletion dramatically reduced fasting-induced lipolysis as indicated by the absence of an increase in plasma free fatty acids in fasted *Atgl*^adipo−/−^(**Figure 1A**). Interestingly, induction by fasting of circulating ketone bodies and plasma FGF21 levels was severely blunted in *Atgl*^adipo−/−^ mice (**Figure 1A**). Next, to characterize changes in hepatic gene expression, we performed transcriptome analyses using microarrays. Principal component analysis (PCA) showed that changes in gene expression between *Atgl*^adipo−/−^ and *Atgl*^adipo+/+^ were predominantly observed in the fasting state (**Figure S1A**). Indeed, among the genes differentially expressed between *Atgl*^adipo−/−^ and *Atgl*^adipo+/+^, 99% were observed in fasted mice while only 0.4% of the genes were differentially expressed in fed mice (**Figure 1B**). Genes expressed at higher levels in response to fasting in *Atgl*^adipo+/+^ compared with *Atgl*^adipo−/−^ mice included genes involved in fatty acid catabolism (*Apoa4*, *Cyp4a14*, *Ehhadh*, *Vnn1*, *Bdh1*) and hepatokines (*Fgf21*, *Inhbe*). Genes with the opposite profile included *Ddit4*, *Lpin1*, and *Sesn1,* which are all p53 target genes (**Figure 1C**). We then performed hierarchical clustering on the differentially expressed genes (DEG). The resulting heatmap clearly discriminated fed from fasted mice and identified 7 clusters showing specific gene expression in response to fasting in the two genotypes (**Figure 1D**). Genes from clusters 4 and 7 were sensitive to fasting but not dependent on *Atgl* expression in adipocytes. However, lack of adipocyte *Atgl* altered the expression of many genes sensitive to fasting. We identified 1182 genes upregulated (cluster 6) and 410 genes downregulated (cluster 2) specifically in fasted *Atgl*^adipo−/−^ compared to *Atgl*^adipo+/+^ mice. Genes from cluster 6 are related to dysregulation in cell cycle and apoptosis and enriched in targets of *Hdac3*, *Creb1*, *Pparg*, *Trp53*, and *Nfkb1* (**Figure 1E**). The fold-changes of the most upregulated genes are shown in supplementary **Figure S1B**. Genes from cluster 2, which were downregulated in the absence of adipocyte ATGL, are mostly involved in metabolism and inflammation. Furthermore, a small cluster of 250 genes (cluster 1) were repressed by fasting, in an adipocyte ATGL-dependent manner. These genes are associated with fatty acid metabolism, steroid metabolism, and circadian rhythm. Accordingly, genes of this cluster are mainly targets of *Arntl*, *Clock*, *Srebf1*, and *Srebf2* (**Figure 1F**). Genes in cluster 5 showed increased expression in response to fasting in *Atgl*^adipo+/+^ but not *Atgl*^adipo−/−^ mice. Gene enrichment analysis revealed that genes of this cluster are mostly involved in fatty acid degradation and peroxisome biogenesis, and enriched in target genes of PPARα (**Figure 1F**). **Figure S1C** lists the genes with the highest fold-changes from this cluster. Accordingly, the expression of well-known PPARα target genes, as assessed by RT-qPCR, was strongly upregulated by fasting in *Atgl*^adipo+/+^ mice but was low or undetectable in the absence of adipocyte ATGL (**Figure 1G**). Similar results were observed for many other PPARα target genes (**Figure S1D**).

**Figure 1:**
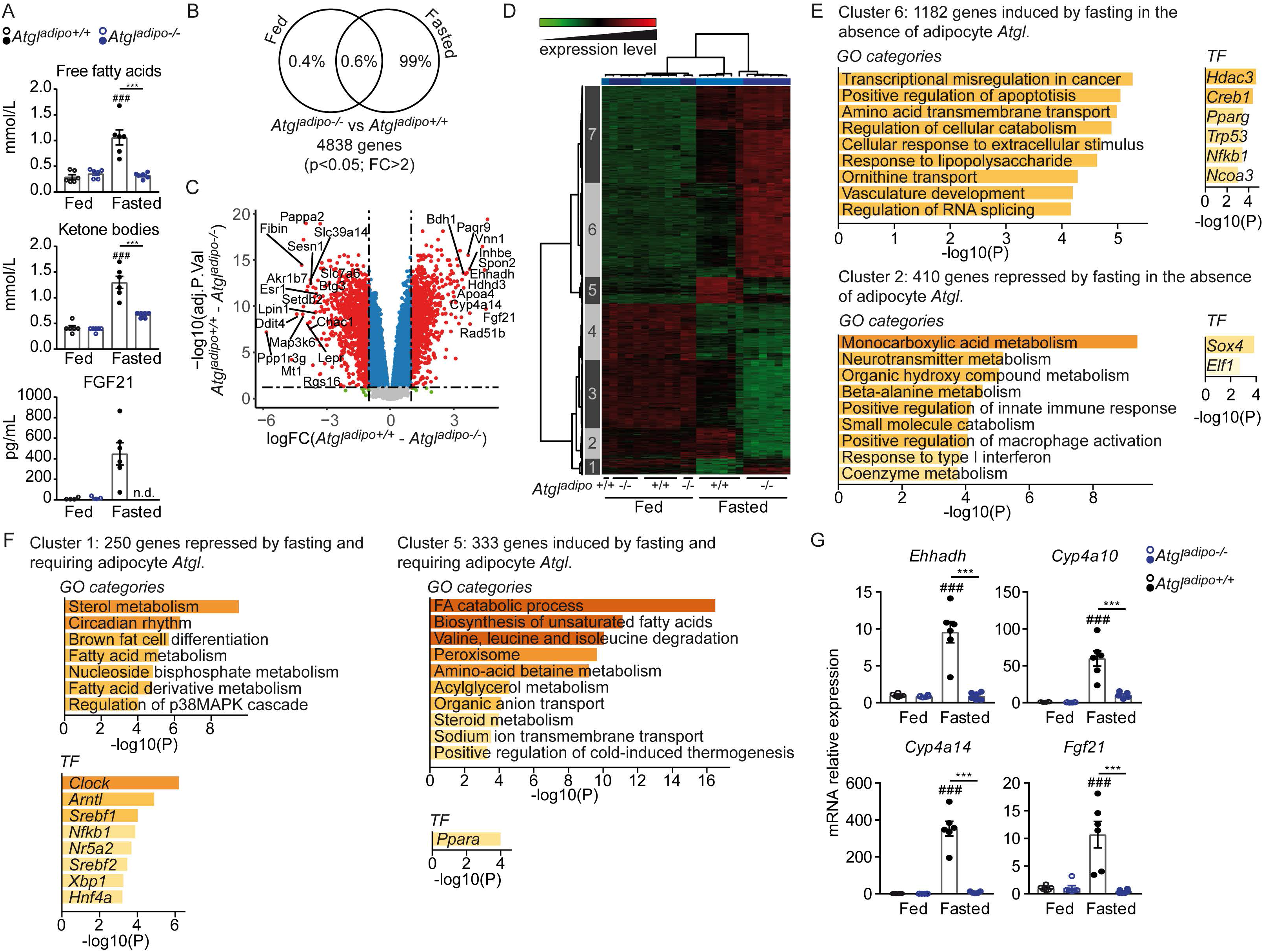
Adipose *Atgl* deficiency altered hepatic gene expression during fasting. (**A–G**) *Atgl*^adipo+/+^ and *Atgl*^adipo−/−^ were fed *ad libitum* or fasted for 24 h. (**A**) Plasma free fatty acid levels and circulating levels of ketone bodies (β-hydroxybutyrate) and FGF21. (**B**) Venn diagram representing the number of genes significantly differentially regulated between *Atgl*^adipo+/+^ and *Atgl*^adipo−/−^ mice in fed and fasted states (FC > 2; *p* < 0.05). (**C**) Volcano plot of differences in gene expression between *Atgl*^adipo+/+^ and *Atgl*^adipo−/−^ in response to fasting. (**D**) Heatmap presenting data from a microarray experiment performed with liver samples (n = 6/group). Hierarchical clustering shows the definition of 7 gene clusters (FC > 2; *p* < 0.05). (**E**) Gene Ontology (GO) analysis and enrichment of transcription factors (TF) of genes specifically up- and downregulated in *Atgl*^adipo−/−^ mice. (**F**) Gene Ontology (GO) analysis and enrichment of transcription factors (TF) of genes specifically up- and downregulated in *Atgl*^adipo+/+^ mice. (**G**) mRNA relative expression of *Ehhadh*, *Cyp4a10*, *Cyp4a14*, and *Fgf21* in the liver measured by qRT-PCR. Results are the mean ± SEM. # fasting effect, * genotype effect, * or # *p* < 0.05, ** or ## *p* < 0.01, *** or ### *p* < 0.001.

Together, these results reveal the important contribution of adipocyte ATGL to the regulation of the liver transcriptome in response to starvation. Further, it strongly suggests a critical role of PPARα in ATGL-dependent hepatic gene regulation during fasting.

### 3.2. Hepatocyte PPARα is critical for the fasting-induced adipose ATGL-dependent effect on liver gene expression

To further delineate the above-identified contribution of hepatocyte PPARα to the fasting-induced adipocyte ATGL-dependent regulation of hepatic gene expression, we compared the liver transcriptomic signature of fasted mice upon adipocyte *Atgl* deletion with that of hepatocyte *Pparα* deletion. We took advantage of our previously published gene expression analysis performed with liver samples from hepatocyte-specific *Pparα* knockout mice that were fasted for 24 h (GEO accession number GSE96559) [3]. Among the 17,508 common hybridized probes (*p* < 0.05) obtained in liver samples in both the PPARα and ATGL experiments, we identified 1,018 probes corresponding to fasting-induced genes that were dependent on ATGL expression in adipocytes, and 443 probes corresponding to fasting-induced genes that were dependent on PPARα expression in hepatocytes (**Figure 2A**). There were 191 probes that shared a common pattern in both experiments (hypergeometric test, *p* = 5.8 × 10^−121^). We also found 1,223 probes and 349 probes corresponding to downregulated genes in fasted *Atgl*^adipo+/+^ and fasted *Pparα*^hep+/+^ mice, respectively, 84 of which were in common (hypergeometric test, *p* = 4.8 × 10^−25^; **Figure 2B**). Hierarchical clustering performed on differentially expressed genes (FC > 2; *p* < 0.05) in the *Pparα* deficiency model allowed the identification of 6 clusters of genes with different patterns. These PPARα-dependent changes in gene expression induced by fasting were analyzed in the *Atgl* adipocyte-deficient model. The resulting heatmap generated from the same gene clustering clearly illustrates the similarity of the gene expression profiles in response to fasting upon adipocyte *Atgl* deletion and hepatocyte *Pparα* deletion, indicating that nearly all genes sensitive to both fasting and hepatocyte PPARα are also dependent on adipocyte ATGL (**Figure 2C**). In contrast, in both heatmaps, genes in clusters 1 and 6 were sensitive to fasting but were not dependent on genotype. Furthermore, genes in cluster 2 were affected by hepatocyte *Pparα* and adipocyte *Atgl* deficiency in both the fed and fasted states; thus, these genes are not specific to fasting. The expression of 281 fasting-induced genes in control *Atgl*^adipo+/+^ and *Pparα*^hep+/+^ mice (cluster 3) was not upregulated in both *Atgl*^adipo−/−^ and *Pparα*^hep−/−^ mice, suggesting that these genes’ expression was dependent on the presence of both ATGL in adipocytes and PPARα in hepatocytes (**Figure 2C**). In accord with these results, we found a strong correlation in the expression level of the genes from cluster 3 (Pearson correlation, R^2^=0.7068680). Among the genes highly dependent on adipocyte ATGL and hepatocyte PPARα, many are well-known target genes of PPARα (*Cyp4a10*, *Cyp4a14*, *Vnn1*, *Ehhadh*, *Acot*s, *Hmgcs2*) (**Figure 2D**). Gene network analysis confirmed the upregulation of genes involved in fatty acid metabolism pathways and enriched in targets of PPARα in response to fasting in control mice only, indicating that both adipocyte ATGL and hepatocyte PPARα are critical for these processes (**Figure 2E**). The 10 genes from cluster 3 with the highest fold-changes are listed in supplementary **Figure S2A**. In contrast, cluster 5 included genes that were upregulated by fasting in the absence of both PPARα in hepatocytes and ATGL in adipocytes (**Figure 2C**). The main network includes genes involved in cell death (**Figure 2E**). **Figure S2B** illustrates the fold-changes of the top 10 most upregulated genes in this cluster. Interestingly, cluster 4 contained a small subset of genes that were upregulated in fasted *Pparα*^hep−/−^ mice but not in *Atgl*^adipo−/−^ mice (**Figure 2C**), suggesting that these genes are constitutively repressed by PPARα. Gene network analysis revealed that these genes are related to the innate immune response (**Figure 2E**). The 10 genes with the highest fold-changes from this cluster are listed in supplementary **Figure S2C**.

**Figure 2:**
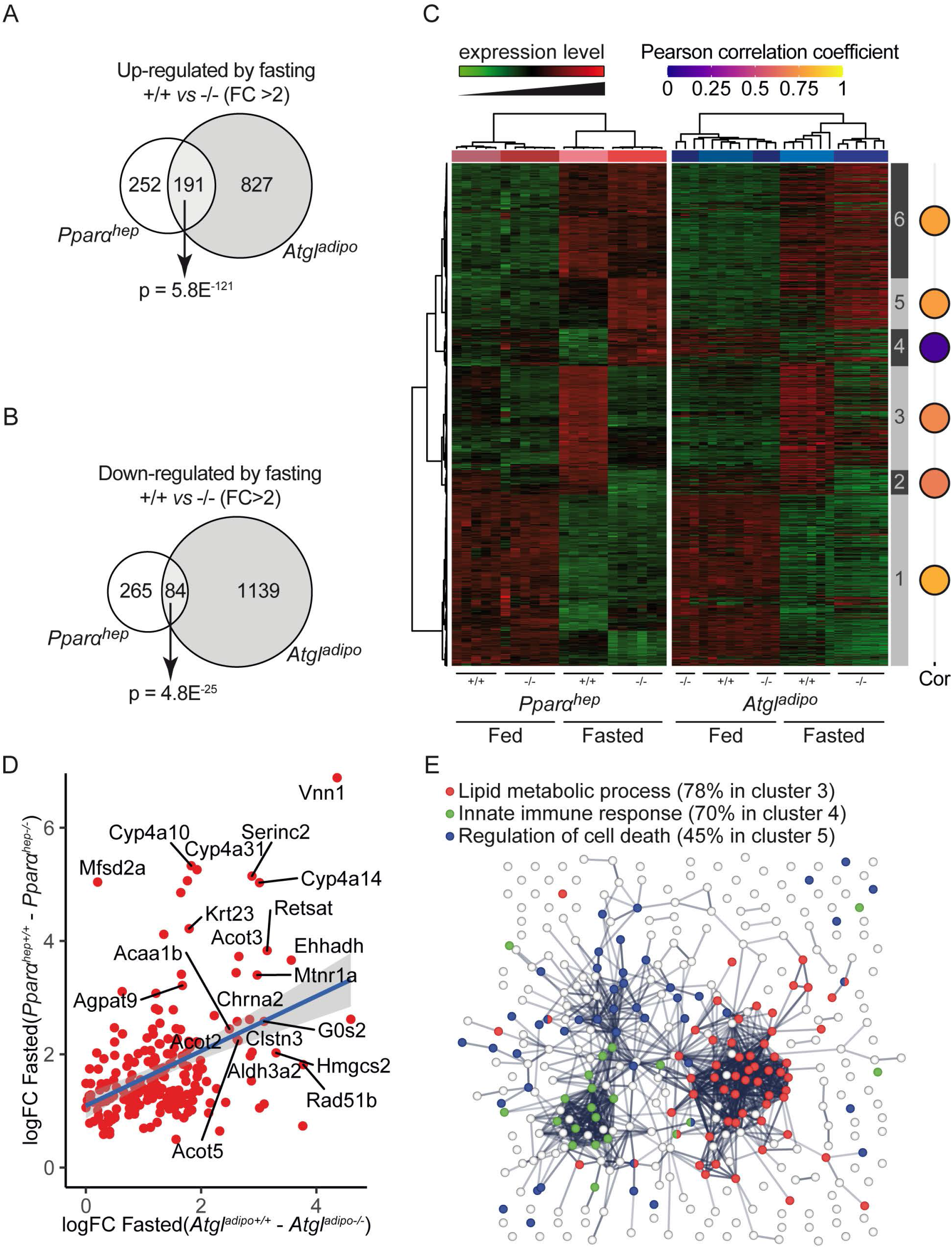
Hepatocyte PPARα is essential for adipose ATGL-dependent hepatic transcriptome in fasted mice. (**A**) Euler diagrams representing the number of probes corresponding to upregulated genes upon fasting and differentially expressed (FC > 2; *p* < 0.05) between *Atgl*^adipo+/+^ and *Atgl*^adipo−/−^ and/or *Pparα*^hep+/+^ and *Pparα*^hep−/−^. (**B**) Euler diagrams representing the number of probes corresponding to downregulated genes upon fasting and differentially expressed (FC > 2; *p* < 0.05) between *Atgl*^adipo+/+^ and *Atgl*^adipo−/−^ and/or *Pparα*^hep+/+^ and *Pparα*^hep−/−^ (**C**) The first heatmap shows data from a microarray experiment performed with liver samples from *Pparα*^hep+/+^ and *Pparα*^hep−/−^ fed *ad libitum* or fasted for 24 h (n = 6/group). Hierarchical clustering shows the definition of 6 gene clusters (FC > 2*; p* < 0.05). The same gene clustering was applied to generate the second heatmap, which represents differentially expressed genes in the liver of *Atgl*^adipo+/+^ and *Atgl*^adipo−/−^ fed *ad libitum* or fasted for 24 h (n = 6/group). Balloon plots on the right represent Pearson correlation coefficients for mean gene expression by experimental group in each cluster of both heatmaps using the *Pparα* experiment as a model. (**D**) Correlation plot of expression levels of genes from cluster 3. (**E**) Gene network analysis of genes from clusters 3, 4, and 5.

Altogether, these results reveal that the PPARα-dependent transcriptomic response in the liver during fasting is highly dependent on adipocyte ATGL, suggesting that adipose-derived fatty acids are critical for hepatocyte PPARα activity.

### 3.3. Acute activation of β_3_-adrenergic signaling is sufficient to induce adipose ATGL-dependent PPARα activity in hepatocytes

To further confirm that adipocyte ATGL-dependent lipolysis is a prerequisite for PPARα activity in the liver, we analyzed the metabolic responses of *Atgl*^adipo−/−^ and *Pparα*^hep−/−^ fed mice following pharmacological induction of adipocyte lipolysis through selective activation of the β_3_-adrenergic receptor using CL316243 (CL). Adipocyte *Atgl* deficiency abolished the increased levels of free fatty acids and circulating ketone bodies and FGF21 in response to CL (**Figure 3A & 3B**). In addition, the hepatic expression of well-described PPARα target genes (*Ehhadh, Cyp4a10*, *Cyp4a14, Fgf21)* in response to CL was dramatically reduced in *Atgl*^adipo−/−^ mice compared with *Atgl*^adipo+/+^ mice, indicating that adipose ATGL is required for CL-induced PPARα transcriptional activity (**Figure 3C**). Hepatocyte *Pparα* deletion did not affect the rise in free fatty acid levels following activation of β_3_-adrenergic signaling (**Figure 3D**). However, similar to what was observed in *Atgl*^adipo−/−^ mice, *Pparα*^hep−/−^ mice exhibited an impaired β_3_-adrenergic response in ketogenesis and FGF21 expression (**Figure 3E & 3F**). We next evaluated the hepatic transcriptome expression patterns in *Pparα*^hep+/+^ and *Pparα*^hep−/−^ mice using microarrays. Differentially expressed genes were subjected to hierarchical clustering. The resulting heatmap highlights marked differences in hepatic gene expression between *Pparα*^hep+/+^ and *Pparα*^hep−/−^ mice in response to CL treatment (**Figure 3G**). We identified 6 clusters of genes with different patterns. Genes in cluster 1 and cluster 4 were regulated by CL but not dependent on hepatocyte *Pparα* expression. Cluster 2 comprises 370 genes that were significantly upregulated following β_3_-adrenergic signaling activation in *Pparα*^hep+/+^ but not in *Pparα*^hep−/−^ mice (**Figure 3H**). These genes are associated with fatty acid metabolism pathways and enriched in targets of PPARα. CL treatment decreased the expression of a small set of genes in a PPARα-dependent manner (cluster 5). Gene ontology analyses revealed that these genes are involved in steroid biosynthesis (**Figure 3H**). The fold-changes of the top 10 up- and downregulated genes from these 2 clusters are presented in **Figure S3A**. Conversely, the expression of some genes changed specifically in response to CL only in the absence of hepatocyte PPARα. Genes from clusters 6 and 3 were respectively more upregulated and downregulated by CL in *Pparα*^hep−/−^ mice than in *Pparα*^hep+/+^ mice. Cluster 6 included 218 genes that are involved in the response to unfolded proteins and regulation of TNF production (**Figure 3I**). Finally, the main pathways repressed by CL in the absence of PPARα in hepatocytes are related to fatty acid degradation and PPAR signaling (cluster 3, **Figure 3I**). The 10 genes with the highest fold-changes are shown in **Figure S3B**. Hepatic transcriptome analysis of CL-treated *Atgl*^adipo+/+^ and *Atgl*^adipo−/−^ also revealed that the absence of adipocyte ATGL had diverse effects on the response to CL (**Figure S4**). The main pathways induced by CL and dependent on adipocyte ATGL were related to fatty acid catabolism and the peroxisome.

**Figure 3:**
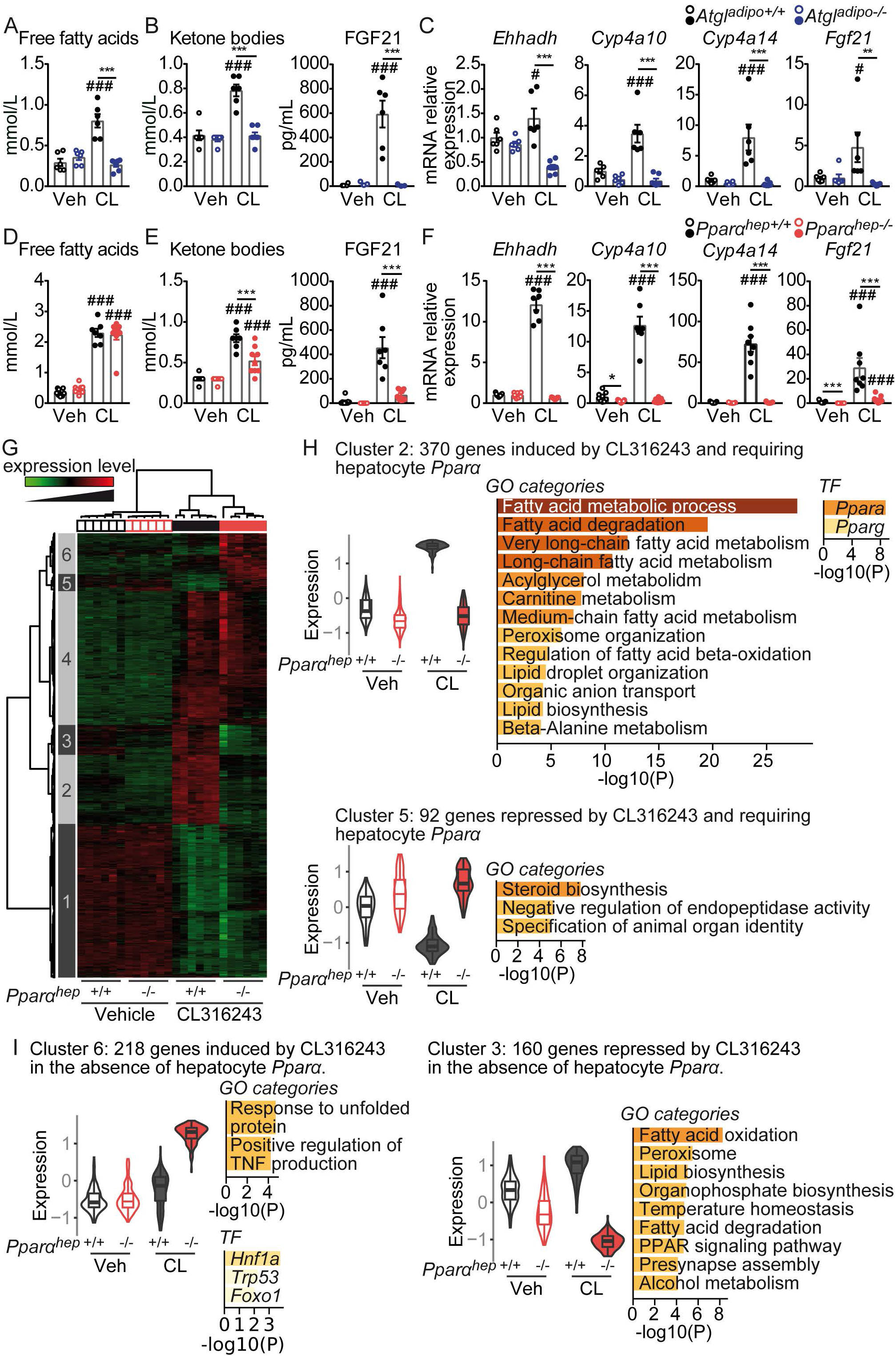
Hepatocyte PPARα activity is induced upon activation of acute β_3_-adrenergic signaling. (**A–C**) *Atgl*^adipo+/+^ and *Atgl*^adipo−/−^ *ad libitum* fed mice received CL316243 at 3 mg/kg body weight or vehicle by gavage and were analyzed 6 hours later. (**A**) Plasma free fatty acid levels. (**B**) Circulating levels of ketone bodies (β-hydroxybutyrate) and FGF21. (**C**) mRNA relative expression of *Ehhadh*, *Cyp4a10*, *Cyp4a14*, and *Fgf21* in the liver measured by qRT-PCR. (**D-I**) *Pparα*^hep+/+^ and *Pparα*^hep−/−^ *ad libitum* fed mice received CL316243 at 3 mg/kg body weight or vehicle by gavage and were analyzed 6 hours later. (**D**) Plasma free fatty acid levels. (**E**) Circulating levels of ketone bodies (β-hydroxybutyrate) and FGF21. (**F**) mRNA relative expression of *Ehhadh*, *Cyp4a10*, *Cyp4a14*, and *Fgf21* in the liver measured by qRT-PCR. (**G**) Heatmap presenting data from a microarray experiment performed with liver samples (n = 6/group). Hierarchical clustering shows the definition of 6 gene clusters (FC > 2; *p* ≤ 0.05). (**H**) Gene expression profile, Gene Ontology (GO) analysis and enrichment of transcription factors (TF) of clusters 2 and 5 (FC > 2; *p* < 0.05). (**I**) Gene expression profile, Gene Ontology (GO) analysis, and enrichment of transcription factors of clusters 3 and 6 (FC > 2; *p* < 0.05).Results are the mean ± SEM. # CL316243 effect, * genotype effect, * or # *p* < 0.05, ** or ## *p* < 0.01, *** or ### *p* < 0.001.

Taken together, these data show that the pharmacological induction of adipocyte lipolysis induced PPARα-dependent responses in the liver even in the fed state. Furthermore, it has been previously shown that inhibition of insulin signaling through mTORC1 is sufficient to induce PPARα activity during fasting [29]. Thus, we tested whether insulin receptor-dependent signaling influences PPARα activity using mice with hepatocyte-specific deletion of the insulin receptor [30,31] (**Figure S5A**). We confirmed that these mice have defective insulin signaling in the liver (**Figure S5B**). The expression of PPARα target genes during fasting was not changed in absence of insulin receptor in hepatocytes (**Figure S5C**). In addition, the deletion of insulin receptor in hepatocytes did not affect the decreased levels of plasma ketone bodies induced by refeeding (**Figure S5D**). Finally, we employed the gain-of-function model of chronically activated insulin signaling, the mouse mutant of hepatocyte-specific inactivation of phosphatase and tensin homolog (PTEN) protein [32,33]. In line with findings in insulin receptor mutant mice, PTEN mutants showed normal ketogenesis during fasting (**Figure S5E**). These results suggest that insulin receptor-dependent signaling in hepatocytes is dispensable for inhibition of hepatocyte PPARα activity in response to feeding. Collectively, these results support a role for fatty acids derived from adipose lipolysis as a determinant signal for hepatocyte PPARα activity.

### 3.4. Hepatocyte PPARα is required for BAT activation and thermogenesis in response to activation of β_3_-adrenergic signaling

β**_3_**-adrenergic signaling not only induces triglyceride lipolysis in WAT but also activates thermogenesis in brown adipocytes. Adipose ATGL is required to maintain mouse body temperature during acute cold exposure, suggesting the importance of lipolysis in providing fatty acids for thermogenesis [34]. To determine the involvement of the WAT-liver axis in BAT thermogenesis, we next investigated the consequences of hepatocyte-specific *Pparα* deletion on BAT activation in response to high-level lipolysis induced by combined pharmacological treatment (CL) and fasting. The increase in plasma free fatty acid levels in response to CL was unaffected by the absence of PPARα in hepatocytes (**Figure 4A**). Interestingly, while blood glucose levels were decreased in both genotypes following CL treatment, the insulin release induced by β**_3_**-adrenergic signaling activation was decreased in *Pparα*^hep−/−^ mice compared with *Pparα*^hep+/+^ mice, suggesting that hepatocyte PPARα is required for insulin secretion (**Figure 4B**). Circulating ketone bodies and plasma FGF21 levels were reduced in fasted *Pparα*^hep−/−^ mice treated with β_3_-adrenergic receptor agonist (**Figure 4C**). In accord with previous results [17], we found that CL decreased the plasma level of free carnitine and enhanced that of medium- and long-chain acylcarnitines (LCACs) levels. In fact, compared with *Pparα*^hep+/+^ mice, *Pparα*^hep−/−^ mice exhibited higher levels of plasma LCACs (**Figure 4D**). CL-induced adipose lipolysis influences the expression of hepatokines. We found that the expression of *Fgf21* and *Inhbe* was increased in response to CL and dependent on hepatocyte PPARα. In contrast, other hepatokine genes such as *Gdf15* and *Fst* were specifically induced in the liver of *Pparα*^hep−/−^ mice following CL treatment (**Figure 4E**). **Figure S6** details the relative expression levels of other hepatokines. We next examined BAT histology. BAT of vehicle-treated mice exhibited many lipid droplets. CL treatment dramatically reduced the number of BAT lipid droplets, indicating BAT activation, but to a lesser extent in *Pparα*^hep−/−^ mice (**Figure 4F**). Expression of genes encoding classic BAT markers was induced upon stimulation of β_3_-adrenergic signaling in control mice. The deletion of hepatocyte *Pparα* led to a specific reduction of *Ucp1* and *Elovl3* mRNA expression in BAT (**Figure 4G**), and UCP-1 protein expression in BAT was lower in *Pparα*^hep−/−^ mice (**Figure 4H**). Finally, we tested adaptive thermogenesis during cold exposure. Acute cold exposure at 4 °C resulted in hypothermia in *Pparα*^hep−/−^ mice compared with *Pparα*^hep+/+^ mice, indicating cold intolerance in the absence of PPARα in hepatocytes (**Figure 4I**). These results revealed that hepatocyte PPARα activation by adipose-derived fatty acids is required for the full activation of BAT. Thus, PPARα activity in hepatocytes mediates the cross-talk between adipose tissues and the liver during lipolysis and thermogenesis.

**Figure 4:**
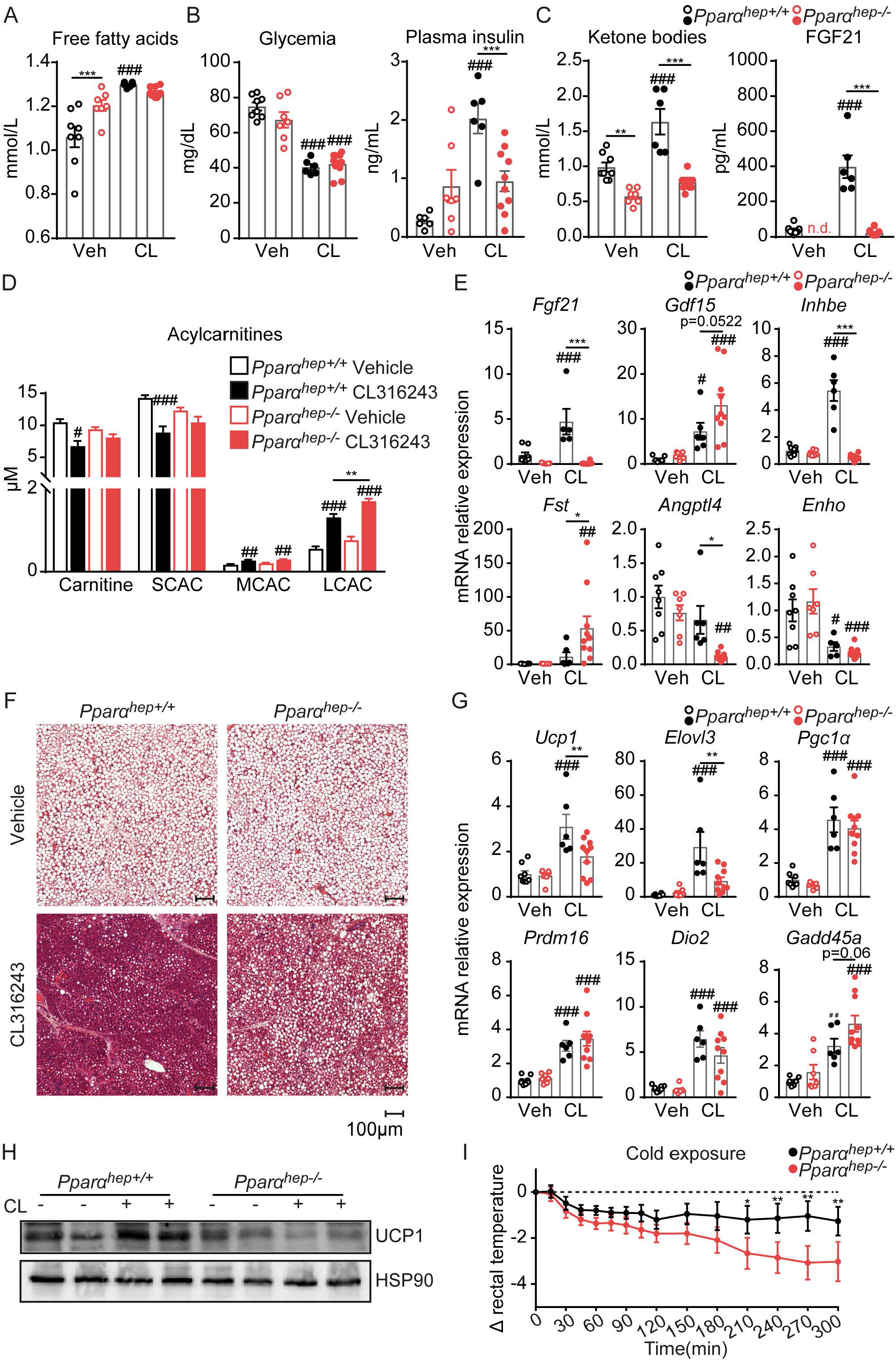
Hepatocyte PPARα deficiency causes reduced BAT activation and defective thermogenesis in response to β_3_-adrenergic signaling activation. (A–H) *Pparα*^hep+/+^ and *Pparα*^hep−/−^ mice were fasted for 10 h, then received CL316243 at 3 mg/kg body weight or vehicle by gavage and were analyzed 6 hours later. (A) Plasma free fatty acid levels. (B) Blood glucose and plasma insulin levels. (C) Circulating levels of ketone bodies (β-hydroxybutyrate) and FGF21. (D) Plasma levels of carnitine, short-chain acylcarnitines (SCAC, C2-C5), medium-chain acylcarnitines (MCAC, C6-C12), and long-chain acylcarnitines (C14-C18). (E) mRNA relative expression of hepatokines in the liver measured by qRT-PCR. (F) Representative histological sections of BAT stained with H&E. (G) mRNA relative expression of *Ucp1*, *Elovl3*, *Pgc1α*, *Dio2*, *Prdm16,* and *Gadd45a* measured in BAT by qRT-PCR. (H) UCP1 protein expression in BAT (the panel shows representative images of all mice in each group). (I) Body temperature after a cold-tolerance test at 4 °C. Results are the mean ± SEM. # CL316243 effect, * genotype effect, * or # *p* < 0.05, ** or ## *p* < 0.01, *** or ### *p* < 0.001.

## 4. Discussion

PPARα is a master regulator of hepatic lipid metabolism during fasting. Consequently, fasted whole-body or liver-specific *Pparα* knockout mice show elevated free fatty acids and defective ketogenesis, and develop hepatic steatosis [2–4]. Although it is generally accepted that fatty acids derived from adipose tissue activate PPARα, a requirement for adipocyte lipolysis for this activation has not been directly addressed [12]. The first and rate-limiting step of adipose triglyceride lipolysis is catalyzed by ATGL. Global and adipocyte-specific *Atgl* deficiency in mice results in defective lipolysis, leading to a shift from lipid to glucose metabolism associated with increased glucose tolerance and insulin sensitivity [18,34]. These mutant mice are also cold intolerant due to defective fatty acid substrates and impaired fatty acid oxidation in cardiac muscle [14]. In the current study, we used mouse models of selective deletion of *Pparα* in hepatocytes and of *Atgl* in adipocytes to elucidate the contribution of ATGL-dependent lipolysis to PPARα activity in the liver. Adipose tissue lipolysis was induced by starvation or by β_3_-adrenergic receptor stimulation. Our data first revealed the strong contribution of adipocyte ATGL to the metabolic response and gene expression in liver during starvation. As previously described [12,18], we found that ATGL-dependent lipolysis is required for fasting-induced ketone body and FGF21 production, two processes under the transcriptional control of PPARα in hepatocytes [2,3,7,8]. In addition, liver gene expression analysis indicated that the genes most sensitive to *Atgl* deficiency included well-established PPARα target genes, such as *Cyp4a10*, *Cyp4a14*, and *Fgf21*. The hepatic transcriptome of fasted mice upon hepatocyte *Pparα* deletion was nearly identical when transposed in the *Atgl* deficiency model, indicating that the expression of genes regulated by hepatocyte PPARα in the liver during fasting also depends on ATGL-dependent lipolysis. These findings indicate that adipocyte ATGL-dependent lipolysis is required for fasting-induced PPARα-dependent hepatic gene expression, and that together they orchestrate the fasting response in the liver. Interestingly, a recent study, using liver-specific deletion of *Atgl* in mice, has reported that hepatic ATGL is not required for the fasting-induced PPARα-dependent responses in the liver, suggesting that adipocyte lipolysis-derived fatty acids are sufficient to activate PPARα independent of the hepatocyte lipolysis [35].

A small subset of genes related to immune response appeared to be upregulated by fasting in the absence of hepatocyte PPARα, but not in the absence of adipocyte ATGL, indicating that these genes are repressed by PPARα independently of the presence of adipocyte-derived ligands. This is in accord with the anti-inflammatory role of hepatocyte PPARα through transrepression which is independent of ligand-induced transcriptional activity [36,37].

Our analysis also revealed that ATGL-dependent lipolysis repressed the hepatic expression of genes that are related to sterol and fatty acid metabolism, and may be involved in the hepatic regulation of sterol regulatory element-binding proteins SREBP-1 and SREBP-2 (cluster 1, Figure 1F). This result is consistent with increased nuclear SREBP-1 levels in hepatocytes of *Atgl*^adipo−/−^ mice after refeeding [38] and suggests that adipose lipolysis–derived fatty acids may suppress SREBP-1 activation in the liver. Previous *in vitro* studies have shown that fatty acids may reduce SREBP-1 activation through the stabilization of the negative regulator insulin-inducible gene 1 (INSIG1) [39,40]. In addition, a recent study reported that PPARα may regulate the processing of the activated form of SREBP-1 during fasting via the upregulation of *Insig2a* gene expression [41]. Furthermore, PPARα can regulate cholesterol levels by reducing the active form of SREBP-2, probably through the increased expression of INSIG proteins [42]. The downregulation of SREBP2-dependent sterol metabolism pathway is in accord with the cholesterol-lowering effects of fibrate PPARα agonists [43]. The down-regulation of SREBP-2 could also explain the down-regulation of SREBP-1 target genes observed in the absence of adipocyte ATGL [44].

Interestingly, we also observed that a large set of genes is regulated by fasting in the absence of adipocyte ATGL, which suggests that these genes are either normally repressed by fatty acids derived from ATGL-dependent lipolysis or specifically induced in the absence of fatty acids as a compensatory mechanism. They are mainly related to cancer and apoptosis and enriched in targets of p53. The tumor suppressor p53 is a transcription factor that regulates the expression of genes controlling cell proliferation and senescence, DNA repair, and apoptosis, and is frequently mutated in hepatocellular carcinoma (HCC) [45]. Free fatty acid levels are elevated in the plasma of HCC patients [46], and lipid metabolic reprogramming has been recently recognized as a hallmark of cancer [47].

We next observed that pharmacological induction of lipolysis with acute CL316243 treatment is sufficient to induce a PPARα-dependent transcriptional response in fed animals. Thus, even in the fed state when insulin is present, the PPARα-dependent responses are induced in *Pparα^hep+/+^* mice and completely abolished in *Pparα^hep−/−^* mice upon adipose lipolysis stimulation. We also found that insulin receptor-dependent insulin signaling does not influence PPARα activity during feeding. In addition, after a fasting period, refeeding is sufficient to decrease PPARα activity even in mice with hepatocyte-specific deletion of the insulin receptor. Although we cannot completely exclude an autonomous negative regulation of PPARα activity by the insulin/mTORC1 axis during feeding [29,48,49], our findings strongly support adipose tissue-derived fatty acids being a dominant signal for hepatocyte PPARα activation. Inhibition of PPARα activity during feeding is likely due to an extrahepatic mechanism, through the inhibition of adipose tissue lipolysis by insulin.

Another interesting finding is the strong influence of hepatocyte PPARα on insulin secretion following β_3_-adrenergic receptor stimulation. It is known that β_3_-adrenergic signaling activation promotes insulin secretion from pancreatic β cells [50]. A recent study reported that this process depends on adipose lipolysis and is essential for the uptake of triglyceride-rich lipoprotein-derived lipids by the BAT and for efficient thermogenesis during β_3_-adrenergic receptor stimulation [15]. Here, we show that hepatocyte PPARα is required for the insulin release, which suggests a cross-talk between the liver and the pancreas that involves PPARα. Whether this effect is mediated by a different composition of plasma lipids or through a hepatic signal, which depends on PPARα activity, remains to be investigated.

Finally, our results reveal for the first time that PPARα activity in hepatocytes is required for full activation of BAT and thermogenesis. Another study first identified the liver as an important tissue for thermogenesis regulation during cold exposure [17]. Simcox et al. showed that acylcarnitines generated by the liver through the fatty acid activation of the nuclear receptor HNF4α in response to cold exposure are required to provide fuel for BAT thermogenesis. Here, we found that activation of hepatic PPARα is also required for BAT activation and cold adaptation. In the current work, we did not observe marked differences in plasma acylcarnitine levels in *Pparα^hep−/−^* mice compared with *Pparα^hep+/+^* mice. Major substrates for BAT activation include glucose [51], fatty acids derived from WAT lipolysis [13,14], circulating triglycerides [15], and acylcarnitines [17]. In our study, however, we could not find any evidence that PPARα influences these parameters. Other PPARα-dependent circulating metabolites include ketone bodies which can act as signaling molecules and whose production is highly dependent on hepatocyte PPARα. However, little is known about the role of ketone bodies in cold adaptation. It has been suggested that ketone bodies may serve as an energy source to fuel thermogenesis [52].

Conversely, we confirmed that hepatic PPARα controls the hepatic expression of FGF21, which could explain the effect on BAT activation as several studies have reported a role for FGF21 in BAT activation [53,54]. Interestingly, we also identified activin E as a novel PPARα-sensitive hepatokine likely to contribute to BAT activation. Activin E is a secreted peptide encoded by the *Inhbe* gene and a member of the transforming growth factor-β family. A recent study reported that activin E is required for cold-induced thermogenesis through the activation of brown adipocytes and induction of beige adipocytes [55]. More studies are needed to identify the hepatic PPARα-dependent signal that triggers BAT activation.

In conclusion, our study supports a dominant role for adipose ATGL in generating tissue-derived fatty acids that trigger hepatocyte PPARα activity, and underscores the critical role of hepatic PPARα not only in the sensing of lipolysis-derived lipids but also in triggering BAT activation. Intact PPARα activity in hepatocytes is required for the cross-talk between adipose tissues and the liver during fat mobilization.

## AUTHOR CONTRIBUTIONS

A.F., G.S., A.P., M.R., C.W., S.S., T.F., Y.L., F.L., V.M., B.T., C.A., A.E., P.D. contributed to experiments, data analysis and provided critical technical support. M.S., P.G., T.L., S.E.S., L.G.P., G.P., E.Z.A., C.P., W.W., N.L., D.L. provided key reagents, intellectual input and reviewed the manuscript. A.F., G.S., A.P., A.M., A.L., H.G. designed experiments, performed experiments, analyzed the data, and wrote the paper.

## ACKNOWLEDGMENTS

We thank all members of the EZOP staff for their help with this project. We thank the GeT-Trix Genotoul facility. We thank Cyrielle Magne-Alibert and Laurent Monbrun from Anexplo for their excellent work on plasma biochemistry. A.F. was supported by a postdoctoral fellowship from Agreenskills, from ANR, and from Région Occitanie. C.A. was supported by a fellowship from the Boulos Foundation and FRM. Additional supported was received by ANRJC “NUTRISENSPIK” (G.P.), by the Leuchtturmprojekt/Flagship Project “Lipases and Lipid Signaling” (A.L.), funded by BioTechMed-Graz, as well as by P28882-B21 (G.S.) and I3535 (A.L.) funded by Austrian Science Fund (FWF). We acknowledge the support of the field of excellence "BioHealth" and NAWI Graz, Austria. This work was funded by ANR (ANR-17-CE14-0015) Hepadialogue (H.G., D.L., C.P., E-Z.A.) and by Région Occitanie (A.M., N.L., D.L., H.G.).

