## Supplementary methods and data for "ATGL-dependent white adipose tissue lipolysis controls hepatocyte PPARα activity"

### Insulin receptor hepatocyte-specific knockout (*IR*$^{hep-/-}$) mice.

Animals carrying LoxP sites flanking the fourth exon of the IR gene (IRlox/lox stock number: 006955; Jackson Laboratory, Bar Harbor, ME, USA) were intercrossed with C57BL/6J mice, which specifically express Cre recombinase in the liver under the transthyretin promoter (TTR-CreTam mice), as previously described [30,31].

At the age of 8 weeks, males IR$^{hep-/-}$ (IR$^{floxed/floxed\ /\ TTR-CreTam+/+}$) and IR$^{hep+/+}$ (IR$^{floxed/floxed\ /\ TTR-CreTam-/-}$) mice received tamoxifen (Tamoxifen Free Base, MP Biomedicals) by intraperitoneal injection (1,5mg/kg) during three consecutive days, to induce deletion of hepatocyte Insulin receptor. Experiments started two weeks after tamoxifen injection.

The IR deletion was confirmed in IR$^{hep-/-}$ mice with PCR and HotStar Taq DNA Polymerase (5 U/μl, Qiagen) using the following primers: IR-forward: 5'-GATGTGCACCCCATGTCTG-3'; IR-reverse: 5'-CTGAATAGCTGAGACCACAG-3' and IRdelta: 5'-GGGTAGGAAACAGGATGG-3'. The amplification conditions were as follows: 95°C for 5 min, followed by 35 cycles of 94°C for 30 sec, 65°C for 30 sec, and 72°C for 45 sec, and a final cycle of 72°C for 7 min. This reaction produced 300-bp, which represented the IR sequence with the floxed allele. The TTR-CreTam allele was detected by PCR using the following primers: forward: 5′-CCTGGAAAATGCTTCTGTCCG-3′ and reverse: 5′-CAGGGTGTTATAAGCAATCCC-3′.

### PTEN hepatocyte-specific knockout (PTEN$^{hep-/-}$) mice.

*Pten*$^{flox/flox}$ mice were mated to *AlbCre* transgenic mice (C57BL6/J background; The Jackson Laboratory, Bar Harbor, Maine, USA), in which expression of Cre is controlled by the promoter of the hepatocyte-specific gene *Albumin,* as previously described [32,33].

### Fasting-refeeding experiment

Twelve-week-old *IR*$^{hep+/+}$ and *IR*$^{hep-/-}$ mice were fasted for 12h (starting at ZT12) or fasted for 12h (from ZT12) and then re-fed until sacrifice (n = 6 mice/genotype/experimental condition).

**Supplementary Table S1. Oligonucleotide sequences for real-time qPCR.**

| Gene | NCBI Refseq | Forward primer | Reverse primer |
|---|---|---|---|
| *Angptl4* | NM_020581.2 | TTTCCCTGCCCTTCTCTACTTG | TACAGGTACCAAACCACCAGCC |
| *Angptl6* | NM_145154.2 | GAATTGCCGCAAACCTCACT | ATGGCCGTCACCTCTCACAG |
| *Angptl8* | NM_001080940.1 | CCCACCAAGAATTTGAGACCTT | ACTGTTGCTGCTCTGCCATCT |
| *Ctsd* | NM_009983 | CTTCGTCCTCCTTCGCGATTAT | GTCCGACGGATAGATGTGAACTT |
| *Cyp4a10* | NM_010011 | TCCAGCAGTTCCCATCACCT | TTGCTTCCCCAGAACCATCT |
| *Cyp4a14* | NM_007822 | TCAGTCTATTTCTGGTGCTGTTC | GAGCTCCTTGTCCTTCAGATGGT |
| *Dio2* | NM_010050.4 | ACAGCTTCCTCCTAGATGCCTACA | GGGAGCATCTTCACCCAGTTT |
| *Ehhadh* | NM_023737 | CGTCTCCTCGGTTGGTGTTC | ATTATCTTCTTTGCAGTATCTAGCTGCTT |
| *Elovl3* | NM_007703 | GCCTCTCATCCTCTGGTCCT | TGCCATAAACTTCCACATCCT |
| *Enho* | NM_027147 | CCGGGCTCAACTCAGGC | TGGCTGTCCTGTCCACACAC |
| *Fetuina* | NM_013465 | ATCGACAAAGTCAAGGTGTGGTCT | TGTCAACTTCCATCTCATACACCACT |
| *Fetuinb* | NM_021564 | CTCGTCAAAGTCACCAAGGCTAT | CACATAGTAAGCAGGGCCAGAC |
| *Fgl1* | NM_145594.2 | TGCAAACCTGAACGGTGTTTAC | TTCAAGGAATACCACCACCCA |
| *Fgf21* | NM_020013.4 | AAAGCCTCTAGGTTTCTTTGCCA | CCTCAGGATCAAAGTGAGGCG |
| *Fst* | NM_008046.2 | TGCTGCTACTCTGCCAGTTCAT | CACTCTTCCTTGCTCAGTTCTGTC |
| *Gadd45a* | NM_007836 | GCGCAGACCCCGGAC | TCCATGTAGCGACTTTCCCG |
| *Gdf15* | NM_011819 | GCTGTCCGGATACTCAGTCCA | TTGACGCGGAGTAGCAGCT |
| *Igf1* | NM_0105124 | GATCTGCCTCTGTGACTTCTTGAA | CAGGTAGAAGAGGTGTGAAGACGA |
| *Igfbp1* | NM_008341.4 | CCTGCCAACGAGAACTCTAT | AGGGATTTTCTTTCCACTCC |
| *Igfbp2* | NM_008342 | GCATGGCCGGTACAACCTTA | GCTGTCCGTTCAGAGACATCTT |
| *Igfbp3* | NM_008343 | CAGGCAGCCTAAGCACCTAC | CTCCTCGGACTCACTGATGTTTC |
| *Inhbe* | NM_008382.3 | TCAGCTTTGCTACCATCATAGACA | CATGGAGCGGTAGGTTGAAGT |
| *Lect2* | NM_010702 | GTGGACAGTACTCTGCTCAAA | TCCCAGTGAATGGTGCATAC |
| *Pgc1α* | NM_008904 | CAATCGGAAATCATATCCAACCA | CTGTGAGGACCGCTAGGAAGT |
| *Pparα* | NM_011144 | CCCTGTTTGTGGCTGCTATAATTT | GGGAAGAGGAAGGTGTCATCTG |
| *Pparα (genotyping)* | NM_011144 | GTACCACTACGGAGTTC | GAATAGTTCGCCGAAAG |
| *Postn* | NM_015784 | GAATGCTGCCCTGGCTATATGA | AATGCCCAGCGTGCCATAAA |
| *Prdm16* | NM_027504.3 | TCCGCGGTCAGCAATAGC | TCACTGCCATCCGACATGTC |
| *Rbp4* | NM_011255 | GCCAAGTTCAAGATGAAGTACTGG | TGTCGTAGTCCGTGTCGATGA |
| *Sepp1* | NM_009155 | AAGATCGCTTACTGTGAGGAGAGG | GCTGAGGTCACAGTTTTACAGAAGTC |
| *Serpinb1a* | NM_025429 | GGACGAGTCCACGGGTCTTA | AGTTTGACGTGGACATCAATGAATTC |
| *Tbp* | NM_013684 | ACTTCGTGCAAGAAATGCTGAA | GCAGTTGTCCGTGGCTCTCT |
| *Ucp1* | NM_009463.3 | CCTGCCTCTCTCGGAAACAA | TGTAGGCTGCCCAATGAACA |

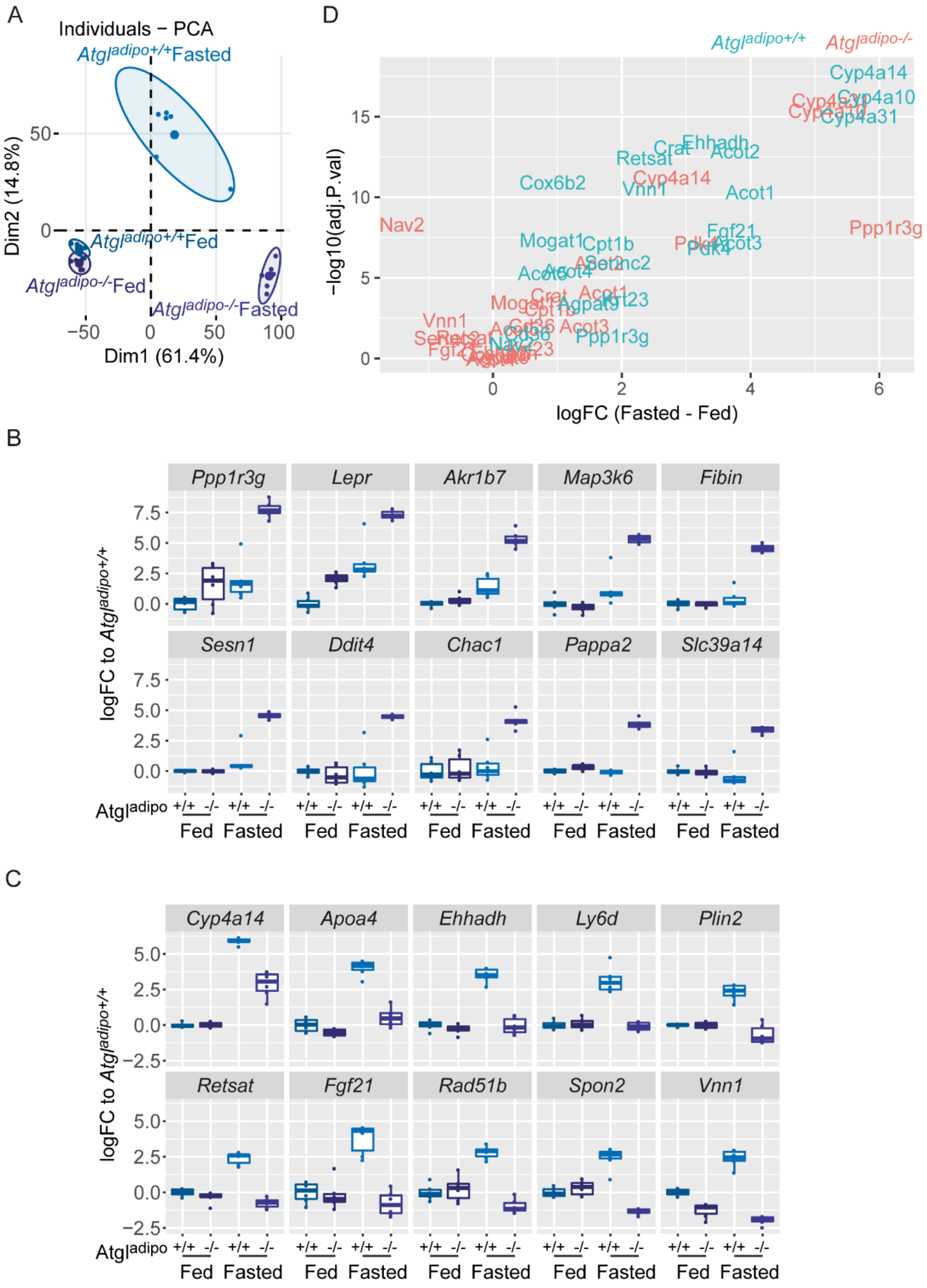


Supplementary figure S1

**Supplementary Figure S1: Adipocyte *Atgl* deficiency altered the fasting-induced hepatic gene expression.** (**A-D**) Microarray experiment performed with liver samples from *Atgl*$^{adipo+/+}$ and *Atgl*$^{adipo-/-}$ mice fed *ad libitum* or fasted for 24 h (n = 6/group). (**A**) Principle component analysis (PCA) score plots of the whole transcriptomic dataset. (**B**) The 10 genes with the highest fold-changes upon fasting in *Atgl*$^{adipo-/-}$ mice. (**C**) The 10 genes with the highest fold-changes upon fasting in *Atgl*$^{adipo+/+}$ mice. (**D**) Volcano plot representing differences in gene expression between *Atgl*$^{adipo+/+}$ and *Atgl*$^{adipo-/-}$ in response to fasting of selected target genes of PPARα.

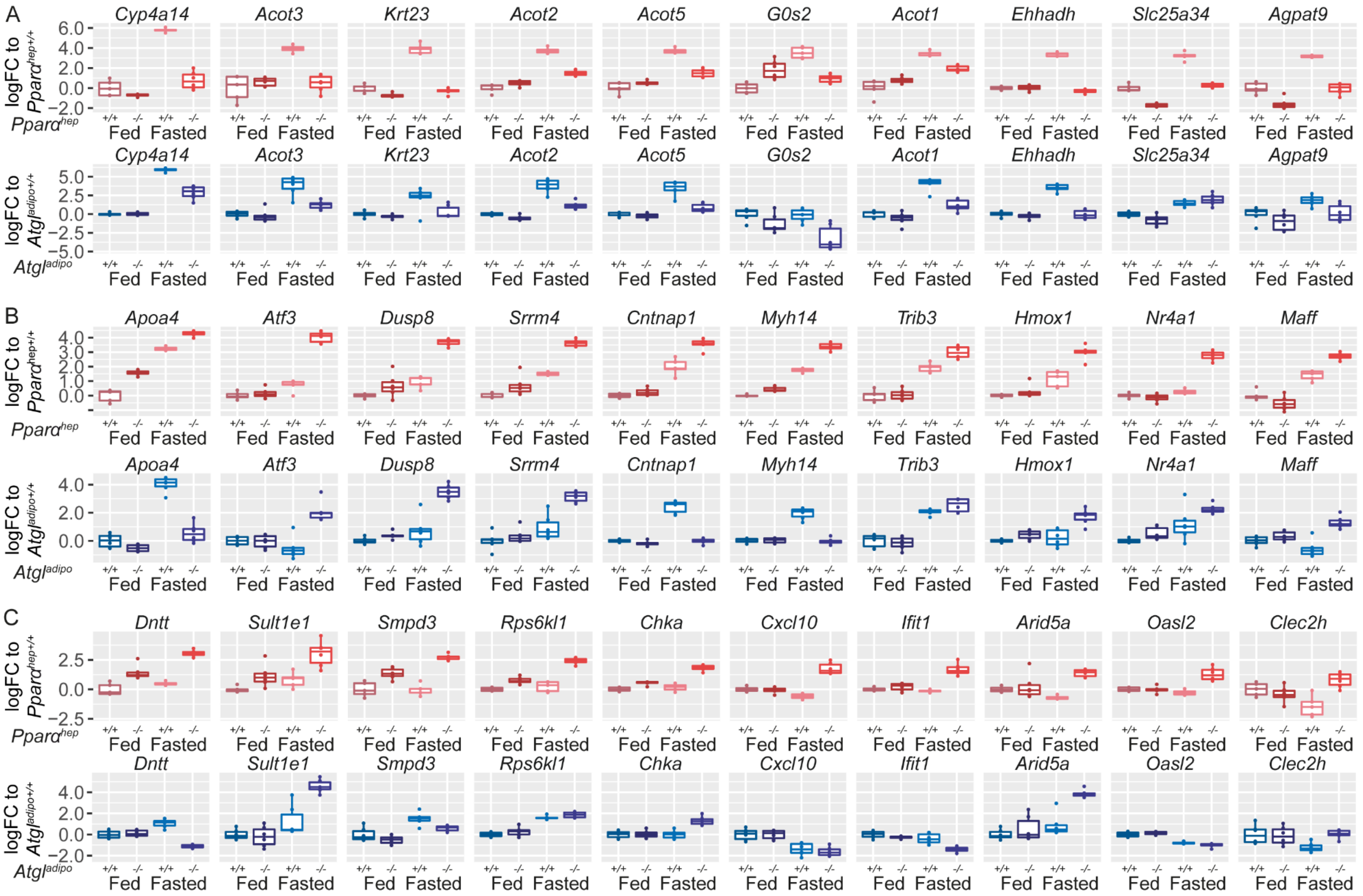


**Supplementary Figure S2: Fasting-induced PPARα-dependent hepatic gene expression analyzed in the *Atgl* deficient model.** (**A-C**) Microarray experiments performed with liver samples from *Atgl*$^{adipo+/+}$ and *Atgl*$^{adipo-/-}$ mice fed *ad libitum* or fasted for 24 h, and from *Pparα*$^{hep+/+}$ and *Pparα*$^{hep-/-}$ mice fed *ad libitum* or fasted for 24 h (n = 6/group). (**A**) The 10 genes from cluster 3 with the highest fold-changes upon fasting in *Pparα*$^{hep+/+}$ mice. Fold-changes for these genes are also shown in the *Atgl* experiment. (**B**) The 10 genes from cluster 5 with the highest fold-changes upon fasting in *Pparα*$^{hep-/-}$ mice. Fold-changes for these genes are also shown in the *Atgl* experiment. (**C**) Fold-changes of the 10 genes most upregulated by fasting only in *Pparα*$^{hep-/-}$ mice, but not in *Atgl*$^{adipo-/-}$ (cluster 4). Fold-changes for these genes are also shown in the *Atgl* experiment.

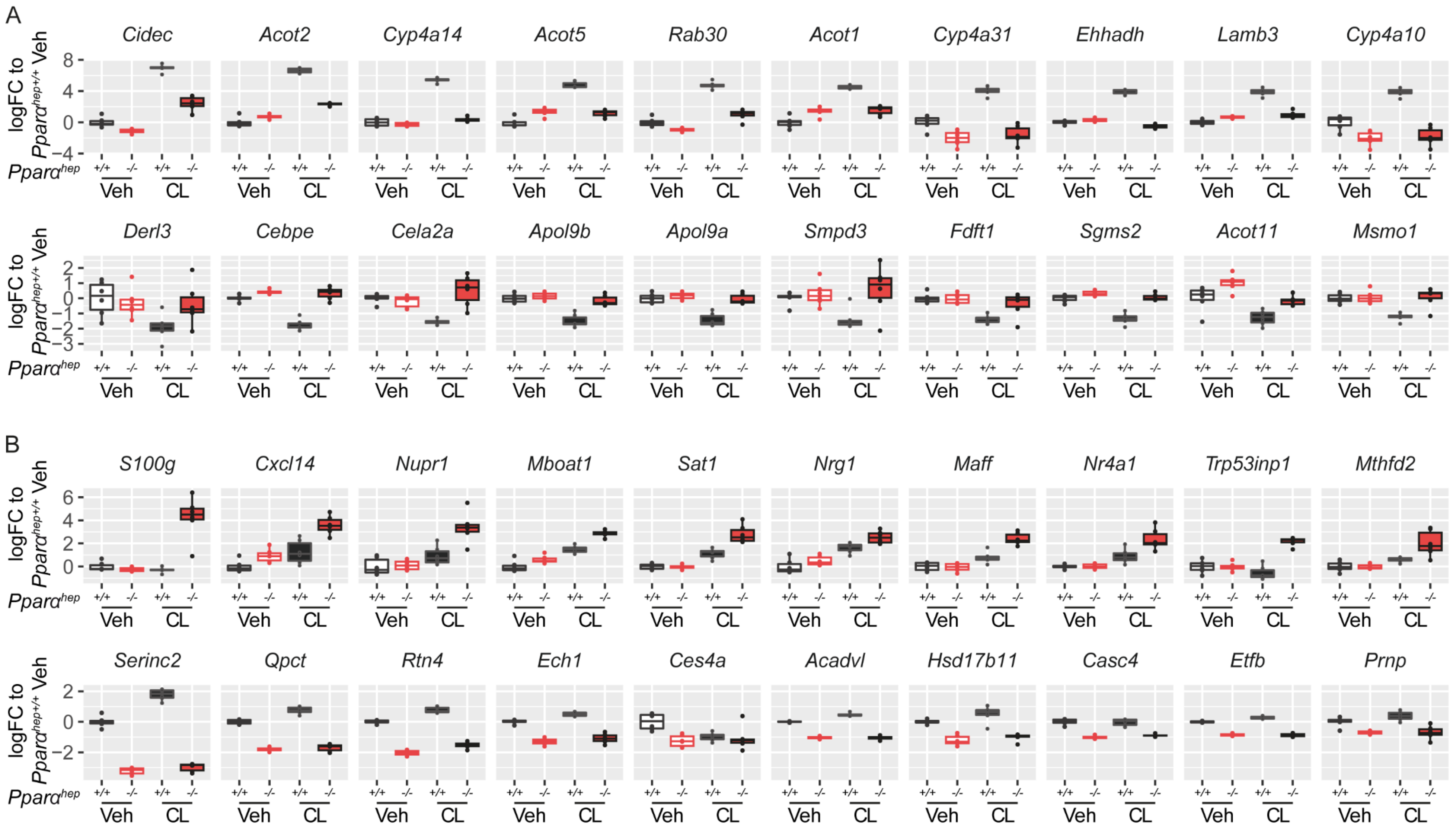


Supplementary figure S3

**Supplementary Figure S3: Hepatocyte *Pparα* deficiency altered hepatic gene expression in response to $\beta_3$-adrenergic receptor stimulation.** (**A-B**) Microarray experiment performed with liver samples from *Pparα*$^{hep+/+}$ and *Pparα*$^{hep-/-}$ fed mice treated with CL316243 (3 mg/kg body weight) or vehicle by gavage for 6 hours (n = 6/group). (**A**) Fold-changes of the 10 genes most upregulated (cluster 2) and of the 10 genes most downregulated by CL316243 (cluster 5) in *Pparα*$^{hep+/+}$ mice. (**B**) Fold-changes of the 10 genes most upregulated (cluster 6) and of the 10 genes most downregulated (cluster 3) by CL316243 in *Pparα*$^{hep-/-}$ mice.

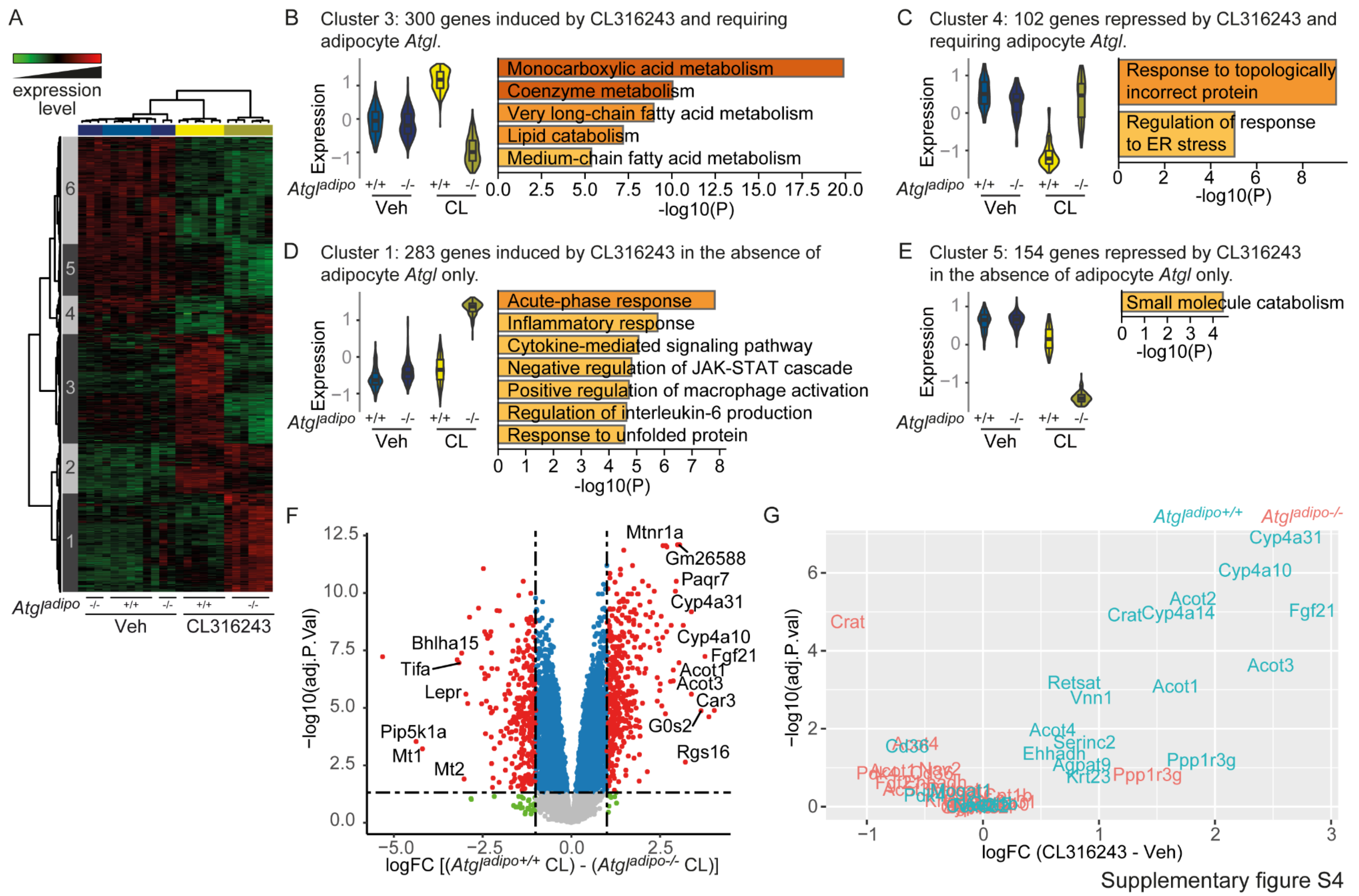


**Supplementary Figure S4: Adipocyte *Atgl* deficiency altered hepatic gene expression in response to $\beta_3$-adrenergic receptor stimulation.** (**A-G**) Microarray experiment performed with liver samples from *Atgl*$^{adipo+/+}$ and *Atgl*$^{adipo-/-}$ fed mice treated with CL316243 (3 mg/kg body weight) or vehicle by gavage for 6 hours (n = 6/group). (**A**) Heatmap with hierarchical clustering shows the definition of 6 gene clusters (FC > 2; $p \leq 0.05$). (**B-E**) Gene expression profile and Gene Ontology (GO) analysis of clusters 3 (**B**), 4 (**C**), 1 (**D**) and 5 (**E**) (FC > 2; $p < 0.05$). (**F**) Volcano plot of differences in gene expression between *Atgl*$^{adipo+/+}$ and *Atgl*$^{adipo-/-}$ in response to CL316243 treatment. (**G**) Volcano plot representing differences in gene expression between *Atgl*$^{adipo+/+}$ and *Atgl*$^{adipo-/-}$ in response to CL316243 treatment of selected target genes of PPARα.

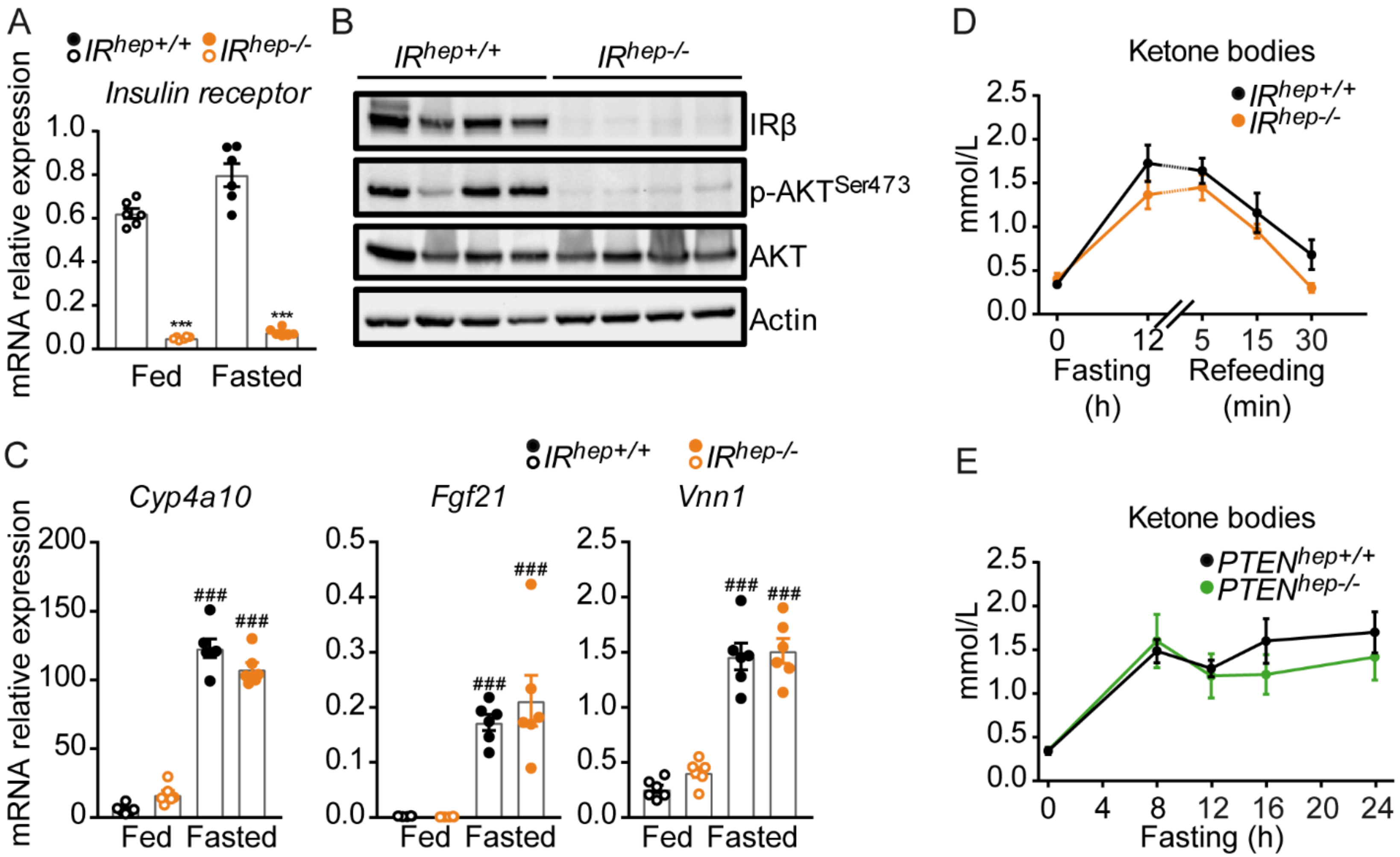


Supplementary figure S5

**Supplementary Figure S5: Modulation of hepatocyte insulin signaling does not influence hepatocyte PPARα-dependent responses in the liver during fasting and refeeding.** (**A**) mRNA relative expression of *Insulin receptor* measured by qRT-PCR in the liver of *IR*$^{hep+/+}$ and *IR*$^{hep-/-}$ mice fed *ad libitum* or fasted for 24 h. (**B**) Western blot analysis of Insulin receptor β (IRβ), phosphorylated AKT, total AKT and β-actin in the liver of *IR*$^{hep+/+}$ and *IR*$^{hep-/-}$ mice fed *ad libitum*. (**C**) mRNA relative expression of *Cyp4a10, Vnn1,* and *Fgf21* measured by qRT-PCR in the liver of *IR*$^{hep+/+}$ and *IR*$^{hep-/-}$ mice fed *ad libitum* or fasted for 24 h. (**D**) Kinetics of circulating levels of ketone bodies (β-hydroxybutyrate) of *IR*$^{hep+/+}$ and *IR*$^{hep-/-}$ mice fasted for 12 h and refed for 30 min. (**E**) Kinetics of circulating levels of ketone bodies (β-hydroxybutyrate) of *PTEN*$^{hep+/+}$ and *PTEN*$^{hep-/-}$ mice fasted for 24 h. Results are the mean ± SEM. # fasting effect, * genotype effect. *** or ### $p < 0.001$.

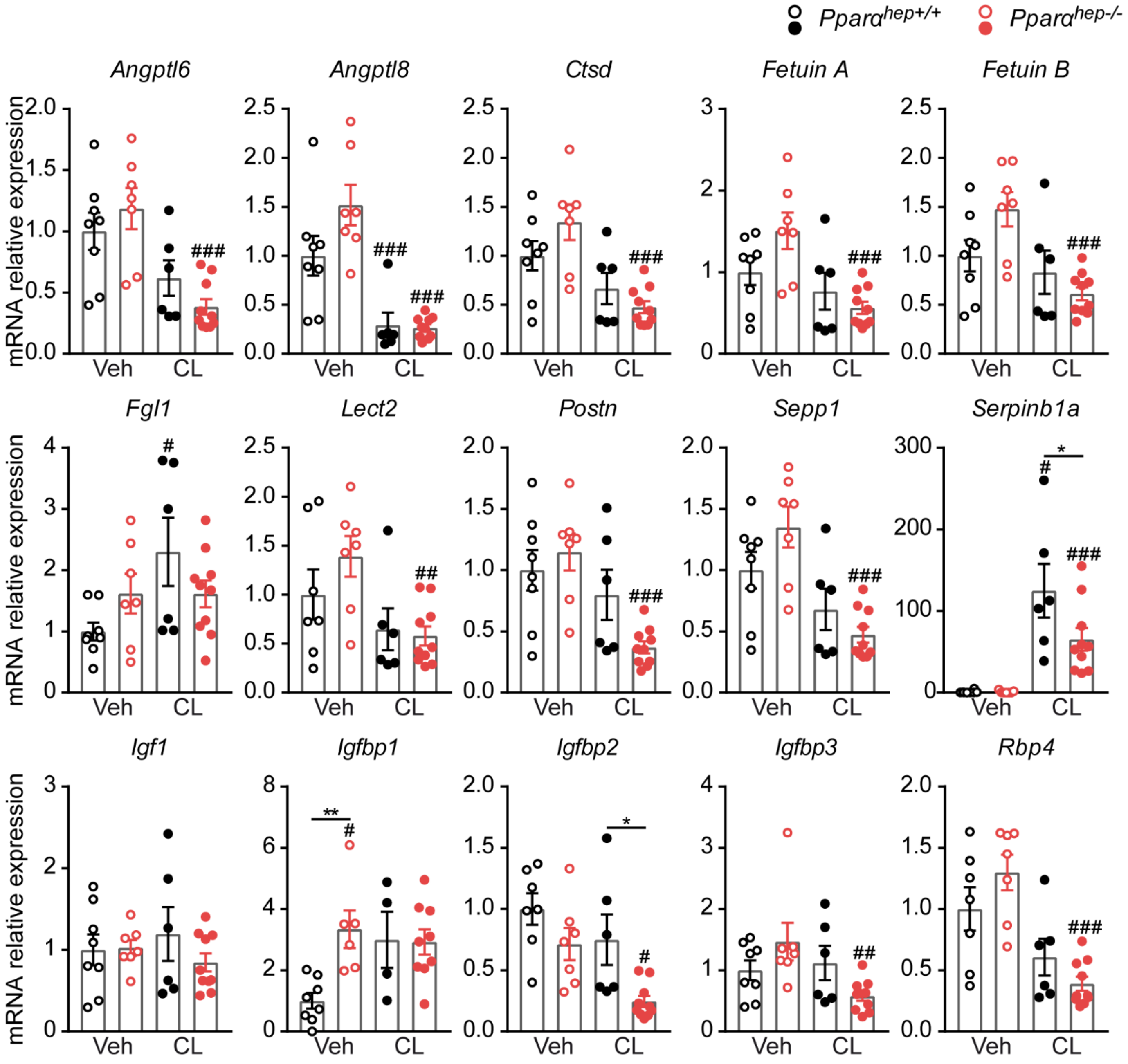


Supplementary figure S6

**Supplementary Figure S6:** Relative gene expression of *Angptl6*, *Angptl8*, *Ctsd*, *Fetuin A*, *Fetuin B*, *Fgl1*, *Lect2*, *Postn*, *Sepp1*, *Serpinb1a*, *Igf1*, *Igfbp1*, *Igfbp2*, *Igfbp3*, and *Rbp4*, as measured by qRT-PCR in fasted *Pparα*$^{hep+/+}$ and *Pparα*$^{hep-/-}$ mice treated with CL316243 (3 mg/kg body weight) or vehicle by gavage for 6 hours. Data are means ± SEM. # CL316243 effect, * genotype effect, * or # $p < 0.05$, ** or ## $p < 0.01$, *** or ### $p < 0.001$.
